# MuteFree: A novel AAV vector system featuring mutation-free ITRs

**DOI:** 10.64898/2026.04.08.717061

**Authors:** Silk J. Shi, Yingjun Lin, Epic Z. Fu, Hermia M. Xu, Robert J. Yang, Yaphia Y. Zhao, Jason Z. Ye, Jack F. Hong, Amy Y. Chen, Xingjian Bai, Bruce T. Lahn

## Abstract

Instability of the inverted terminal repeats (ITRs) in AAV transfer plasmids has long hindered consistent and efficient production of therapeutic AAV vectors. The palindromic, GC-rich ITR sequence readily forms secondary structures, making them highly mutable in transfer plasmids. Indeed, a recent survey observed mutated ITRs in ∼40% of AAV transfer plasmids from labs around the world. Conventional strategies to mitigate this issue – such as using specialized *E. coli* strains, suboptimal culture conditions, or modified ITR sequences – have limited effect and often compromise plasmid and AAV yield. Here, by combinatorial optimization of the plasmid backbone structure and ITR flanking sequences, we established MuteFree, an AAV transfer plasmid system that eliminated ITR mutations for both single-stranded AAV (ssAAV) and self-complementary AAV (scAAV). Specifically, MuteFree reduced ITR mutation rates from a range of 32-100% in various transfer plasmids tested to 0% after serial passage of host *E. coli* for >160 population doublings. Moreover, in three GMP-grade AAV plasmid manufacturing projects initially cancelled due to severe and incurable ITR mutations, replacing conventional backbone with MuteFree completely solved the problem, reducing mutation occurrence to zero under standard GMP manufacturing conditions. Notably, MuteFree supports the packaging of potent AAV virus. The MuteFree system thus presents a robust solution to ITR instability, enabling high-fidelity and high-yield AAV production of AAV-based gene therapy vectors that is fully compatible with existing GMP manufacturing workflows.

## INTRODUCTION

Recombinant adeno-associated virus (AAV) is widely used as a gene delivery vector in gene therapy applications due to its strong in vivo infectivity, broad and tunable tropism, favorable safety profile, and low immunogenicity (1,2). Wildtype AAV is replication-defective by itself, and can only replicate in cells coinfected with certain helper viruses such as adenovirus and herpesviruses (3). Structurally, wildtype AAV contains a ∼4.7 kb single-stranded linear DNA genome encapsulated in an icosahedral protein capsid (4). The viral genome carries only two genes, rep which encodes several alternatively transcribed and spliced variants of DNA polymerase involved in viral genome replication and packaging, and cap which encodes several alternatively transcribed and spliced variants of the capsid protein involved in viral encapsulation (5). The rep and cap genes are flanked by two inverted terminal repeats (ITRs) at the two ends of the genome that act as *cis*-acting packaging signals essential for viral genome replication and encapsulation during virus packaging, and long-term viral genome maintenance in infected cells (5). For recombinant AAV vectors used in gene delivery, the viral genes are replaced with one or more therapeutic transgenes, leaving the flanking ITRs as the only leftover viral genome sequence delivered into target cells along with the transgenes.

To produce recombinant AAV, cultured packaging cells (typically 293 or 293T) are transfected with three plasmids, including 1) transfer plasmid that carries the transgene of interest flanked by two ITRs, 2) rep-cap plasmid that expresses the rep and cap genes, and 3) helper plasmid that expresses certain adenovirus genes that promote AAV packaging by modulating rep and cap function as well as inhibiting host cell defense (6,7).

The ITRs contain a partially palindromic, GC-rich (∼70%) sequence that form a T-shaped secondary structure in the AAV genome. On the transfer plasmid, this complex sequence makes the ITR regions highly unstable during plasmid propagation (8). Recently, we surveyed the integrity of ITRs in hundreds of AAV transfer vectors sourced from academic and industrial labs worldwide, and found that nearly 40% of them exhibited ITR mutations (9). Similarly, surveys of the most widely used AAV plasmids deposited in Addgene revealed that nearly 70% of their ITRs have mutated (9). In this study, quantitative analysis of conventional single-stranded AAV (ssAAV) or self-complementary AAV (scAAV) transfer plasmids showed as high as 100% ITR mutations after a stringent passage test.

Currently, seven gene therapy drugs using AAV vectors have been approved by the FDA, and there are hundreds of AAV-based gene therapy drug candidates in preclinical or clinical development (10). A common challenge for all of them is the manufacturing of high-quality, high-yield AAV vectors. Capsids containing no or truncated genomes often make up a large fraction of the total capsids during manufacturing, yet they deliver no functional genes and may contribute to unnecessary inflammatory response (11). The high rate of ITR mutations greatly exacerbates this problem, as defective ITRs can reduce packaging yield, increase the fraction of capsids containing no or truncated genomes, and compromise the functional efficacy of the final therapeutic virus (12,13).

In this study, we systematically redesigned the backbone of AAV transfer vectors, resulting in the MuteFree system that reduced ITR mutation rate to below detection limit for both ssAAV and scAAV. We further show that MuteFree supports the packaging of potent AAV virus, including when production is performed under standard GMP manufacturing conditions. MuteFree thus represents a robust system for high-fidelity and high-yield manufacturing of therapeutic AAV vectors.

## RESULTS

### Optimization of ITR-adjacent sequence improves ITR stability

The stability of ITRs on AAV transfer plasmids was tested in ten passages. A single-clone *E. coli* culture harboring a validated AAV transfer plasmid was inoculated into 100 mL of LB medium containing the appropriate selective antibiotics. After overnight incubation at 37°C, 1 μL of the culture was transferred into 100 mL of fresh LB medium. The same incubation and subculturing were repeated for a total of ten passages, constituting ∼166 doublings from the original culture. At this point, the culture was plated, and ten single colonies were isolated. To evaluate the potential effects of plasmid DNA extraction and transformation on ITR stability, minipreps were performed four times, and each plasmid preparation was independently retransformed into competent *E. coli*. From each transformation, four single colonies were collected. All the above colonies were subjected to ITR sequence validation to evaluate their stability. **Figure 1** illustrates the process.

**Figure 1.**
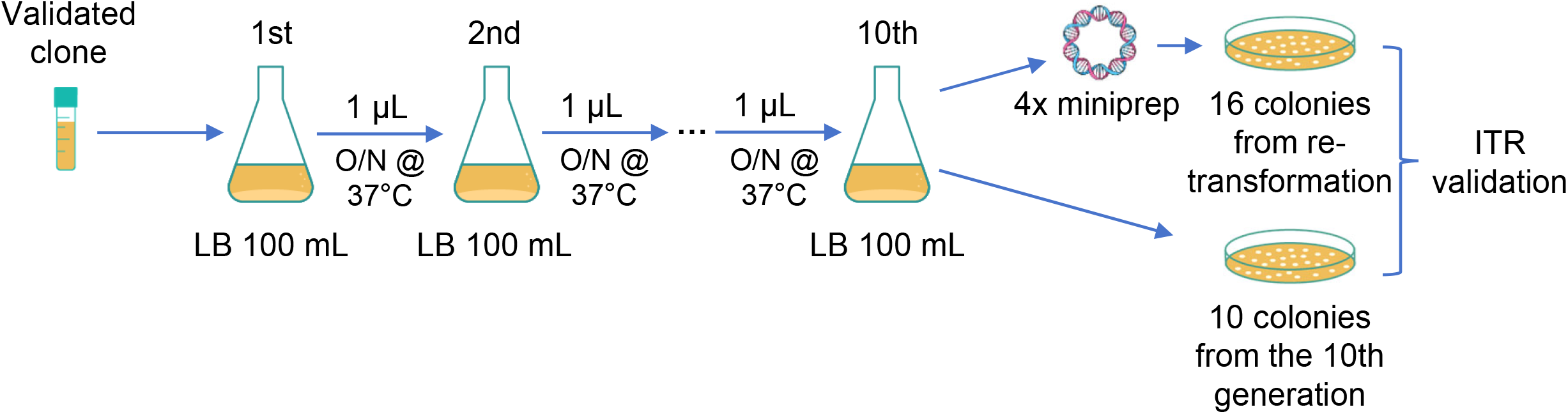
Workflow for assessing AAV ITR stability. A single *E. coli* clone containing a sequence-validated AAV transfer plasmid was serially passaged for ten generations. Plasmid DNA from the tenth generation was isolated in four independent preparations, each retransformed into competent *E. coli*. Four colonies from each retransformation (16 total) and ten colonies from the 10th generation culture were analyzed for ITR integrity by Sanger sequencing.

The baseline ssAAV construct, ssAAV1.0 (**Figure 2A**), contains two 130-nt ITRs flanked by an 11-nt GC-rich region (CCTGCAGGCAG, 73% GC). The plasmid has a conventional backbone featuring an ampicillin resistance gene and a pUC origin of replication (Ori) located 72 nt upstream of the 5′ ITR. The vector carries an expression cassette of CMV promoter driving EGFP as the transgene payload. Based on global plasmid data and available Addgene vector maps, AAV transfer vectors with this backbone and ITR configuration represent the most widely used design in both academic and industrial settings (9).

**Figure 2.**
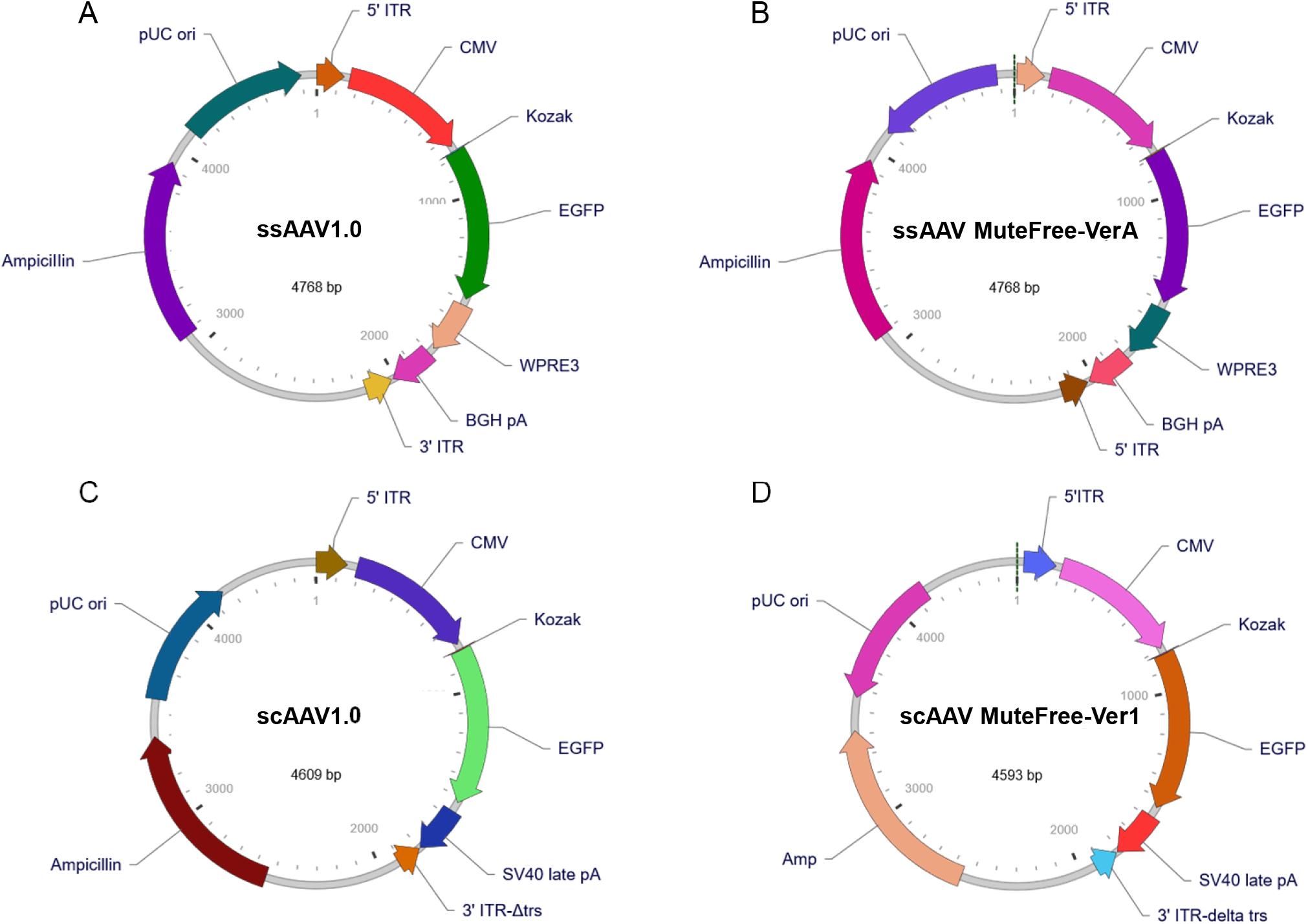
Vector maps for (**A**) ssAAV1.0 (baseline vector), (**B**) ssAAV MuteFree-VerA, (**C**) scAAV1.0 (baseline vector), and (**D**) scAAV MuteFree-Ver1.

Stability test for ssAAV1.0 was conducted twice. Both ITRs from 20 single colonies after ten serial passages were examined by Sanger sequencing. Of these, 50.0% (10/20) of the 5’ ITR showed mutations, including depletion of four bases in A-A’ region and complete loss of the B-B’/C-C’ arms, whereas no mutations were detected in the 3’ ITR (**Table 1**). As for the single colonies after retransformation, ITRs in 32 colonies were fully sequenced, revealing 46.9% (15/32) mutation in 5’ ITR and 0% (0/32) in 3’ ITR, a pattern similar to that observed in the tenth-passage clones (**Table 1**). Collectively, the total 5’ ITR mutation rate of 48.1% (25/52) demonstrate that the 5′ ITR in the commonly used AAV transfer vector exhibits substantial instability (**Table 1**). Furthermore, the mutation rates before and after retransformation were comparable (50.0% vs 46.9%), indicating that plasmid extraction and transformation procedures have minimal impact on ITR mutation. The stability of ITR in all subsequent vectors was evaluated based on the same ten serial passages. The overall ITR stability was determined by combining the data from all clones before and after retransformation.

**Table 1.**
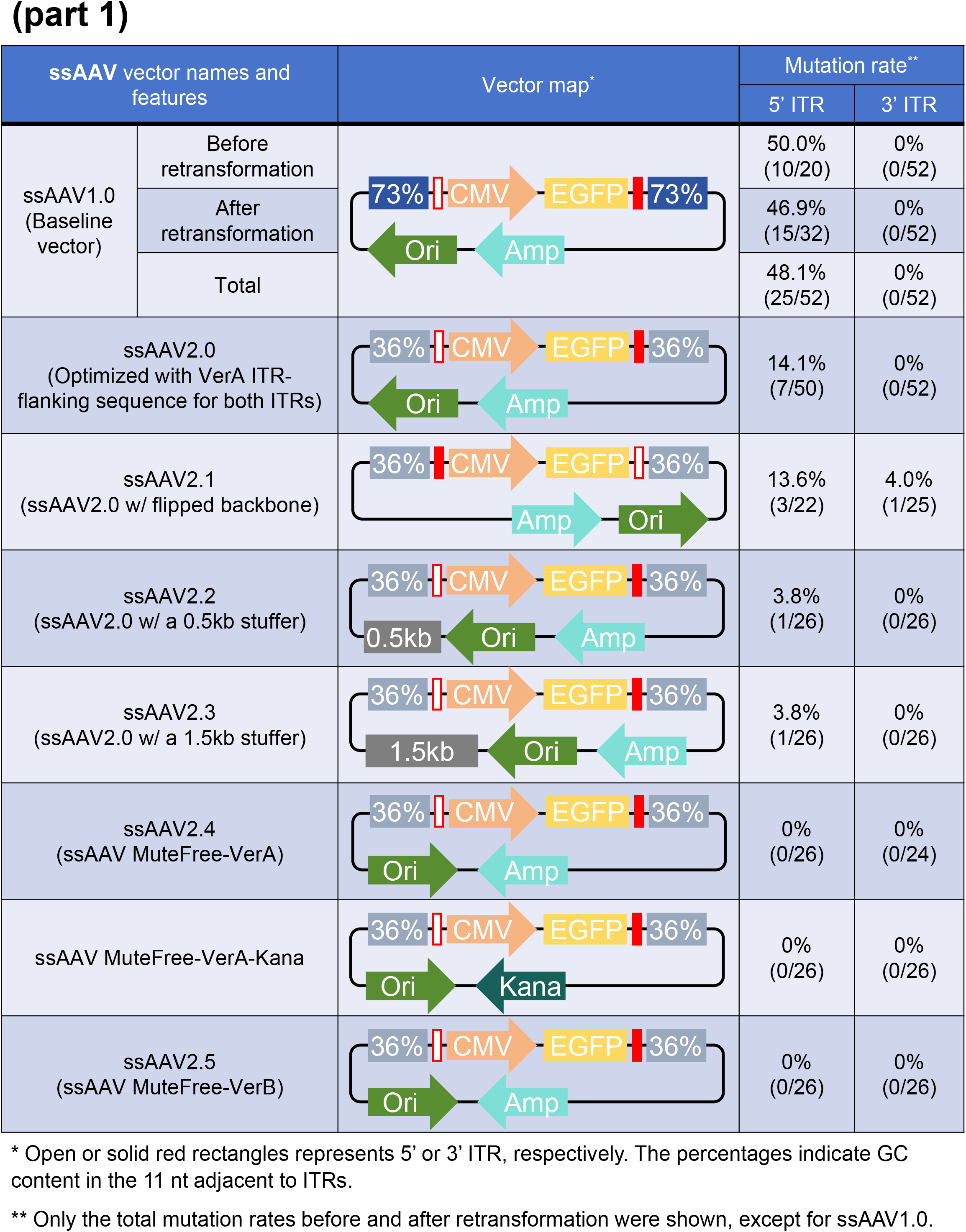
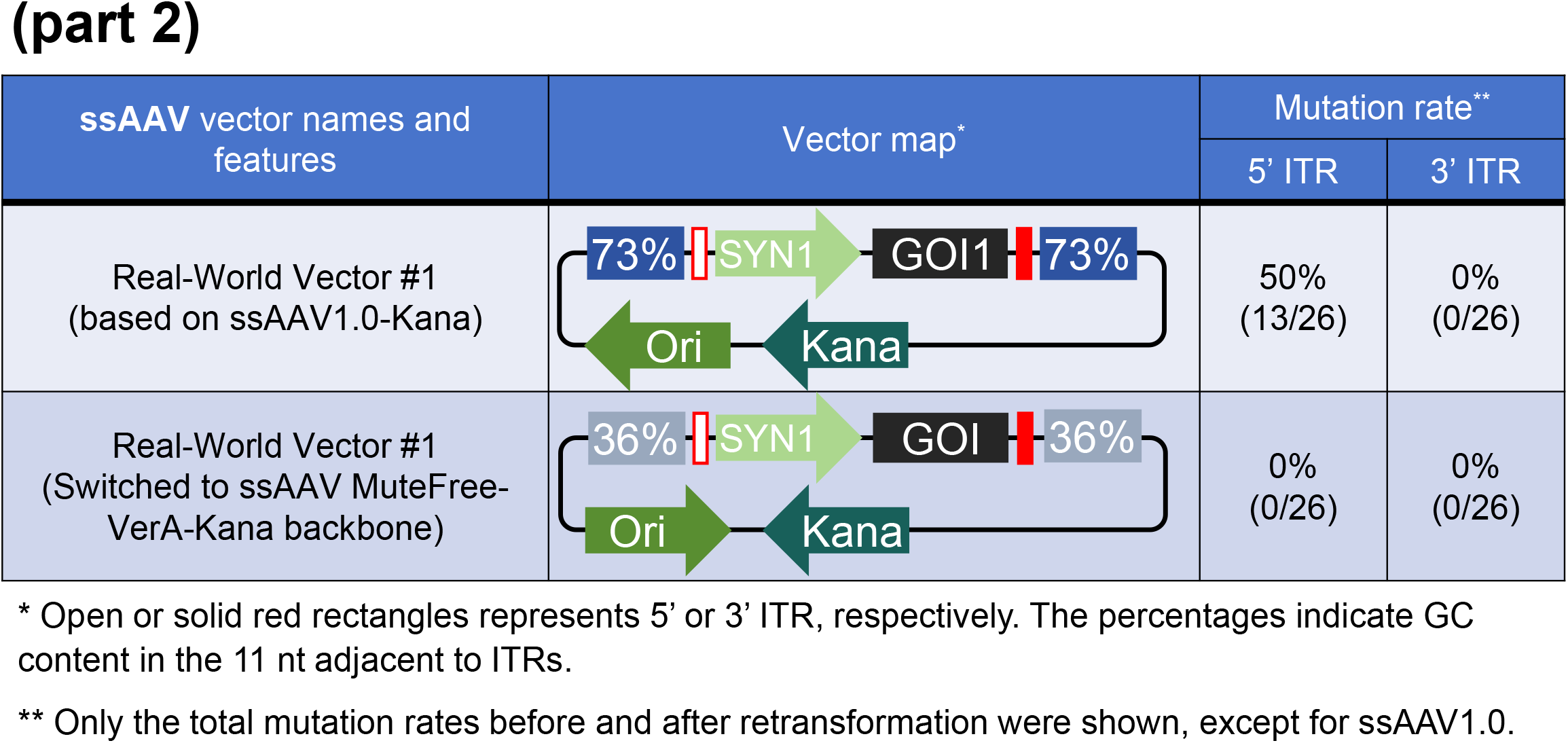
Features of ssAAV transfer vectors and their ITR mutation rate.

| ssAAV vector names and features |  | Vector map* | Mutation rate** |  |
| --- | --- | --- | --- | --- |
|  |  |  | 5' ITR | 3' ITR |
| ssAAV1.0<br>(Baseline vector) | Before retransformation |  | 50.0%<br>(10/20) | 0%<br>(0/52) |
|  | After retransformation |  | 46.9%<br>(15/32) | 0%<br>(0/52) |
|  | Total |  | 48.1%<br>(25/52) | 0%<br>(0/52) |
| ssAAV2.0<br>(Optimized with VerA ITR-flanking sequence for both ITRs) |  |  | 14.1%<br>(7/50) | 0%<br>(0/52) |
| ssAAV2.1<br>(ssAAV2.0 w/ flipped backbone) |  |  | 13.6%<br>(3/22) | 4.0%<br>(1/25) |
| ssAAV2.2<br>(ssAAV2.0 w/ a 0.5kb stuffer) |  |  | 3.8%<br>(1/26) | 0%<br>(0/26) |
| ssAAV2.3<br>(ssAAV2.0 w/ a 1.5kb stuffer) |  |  | 3.8%<br>(1/26) | 0%<br>(0/26) |
| ssAAV2.4<br>(ssAAV MuteFree-VerA) |  |  | 0%<br>(0/26) | 0%<br>(0/24) |
| ssAAV MuteFree-VerA-Kana |  |  | 0%<br>(0/26) | 0%<br>(0/26) |
| ssAAV2.5<br>(ssAAV MuteFree-VerB) |  |  | 0%<br>(0/26) | 0%<br>(0/26) |

\* Open or solid red rectangles represents 5' or 3' ITR, respectively. The percentages indicate GC content in the 11 nt adjacent to ITRs.
\*\* Only the total mutation rates before and after retransformation were shown, except for ssAAV1.0.

Table 1 (part 2)
| ssAAV vector names and features | Vector map* | Mutation rate** |  |
| --- | --- | --- | --- |
|  |  | 5' ITR | 3' ITR |
| Real-World Vector #1<br>(based on ssAAV1.0-Kana) |  | 50%<br>(13/26) | 0%<br>(0/26) |
| Real-World Vector #1<br>(Switched to ssAAV MuteFree-VerA-Kana backbone) |  | 0%<br>(0/26) | 0%<br>(0/26) |

\* Open or solid red rectangles represents 5' or 3' ITR, respectively. The percentages indicate GC content in the 11 nt adjacent to ITRs.
\*\* Only the total mutation rates before and after retransformation were shown, except for ssAAV1.0.

Given previous findings suggesting that ITR stability can be influenced by adjacent sequences (9), we next examined a series of ssAAV1.0 derivative plasmids in which the 11 ITR-flanking nucleotides were systematically modified to a large number of alternative sequences. These variants spanned a range of GC contents from 0% to 64%, with multiple sequence variations tested at each GC level. After subjecting them to the same ITR stability test, two optimal one, named version A (VerA) and version B (VerB), both containing 36% GC, were chosen for further analysis. One plasmid we tested was ssAAV2.0, which was identical to ssAAV1.0 except incorporating the VerA ITR-adjacent sequence for both ITRs. It lowered the 5’ ITR mutation rate from the previous 48.1% (25/52) in ssAAV1.0 to 14.1% (7/50) and maintained 0% (0/52) mutation rate for the 3’ ITR, while preserving structural integrity of the plasmid (**Table 1**). The plasmid for which the ITR-adjacent sequence was replaced with VerB performed equally well.

### MuteFree system eliminates ITR mutations in ssAAV

While optimizing the flanking sequences reduced the ITR mutation rate to 14.1%, it did not fully solve the problem. We then moved on to further improve ITR stability by optimizing the backbone. To investigate the effect of backbone structure on ITR stability, a series of vectors were constructed by systematically altering the placement and spacing of each functional element within the ssAAV2.0 backbone. The effect of these backbone configurations on ITR stability was assessed using the abovementioned serial passage assay. While most of them resulted in only modest changes, several variants exhibited notable effects and warrant further discussion. We first constructed ssAAV2.1 by inverting the entire backbone of ssAAV2.0. The 5’ and 3’ ITR mutation rates in ssAAV2.1 were 13.6% (3/22) and 4.0% (1/25), respectively. These mutation rates were comparable to those observed for ssAAV2.0 (**Table 1**). It is worth noting that we define the ITR located proximal to the Ori as 5’ ITR; thus, relative to the genetic cargo, the 5’ and 3’ ITR in ssAAV2.1 were switched compared to ssAAV2.0. The 5’ ITR in both cases were significantly less stable compared to the 3’ ITR, suggesting that backbone configuration indeed affects ITR stability and highly biased instability of 5’ ITR is more influenced by backbone configuration rather than by the genetic cargo within the ITRs.

By rearranging the positions and spacing of the various backbone components in ssAAV2.0 while maintaining the ITR-adjacent sequence, we tested a large number of vector configurations for their ITR stability. The majority of them didn’t improve ITR stability. But we did identify three backbone configurations that dramatically improved ITR stability. Two backbone variants were generated by inserting an insulator sequence of 0.5 or 1.5 kb inserted between the Ori and the 5′ ITR in ssAAV2.0 (named ssAAV2.2 and ssAAV2.3, respectively). These modifications reduced 5’ ITR mutation rate to 3.8% (1/26) (**Table 1**). However, the increased plasmid length introduced by the insulator sequences may compromise transfection efficiency during vector packaging, potentially reducing AAV yield. In addition, incorporation of nonessential sequences into an AAV transfer plasmid may pose challenges for regulatory compliance, as these elements would require comprehensive safety evaluation prior to clinical application. The third vector, ssAAV2.4, showed no detectable mutations in either the 5′ or 3′ ITRs (0/26) without introducing additional sequences (**Table 1**). We thus considered ssAAV2.4 to be the optimal solution when considering both transfection efficiency and clinical regulatory requirements. Therefore, ssAAV2.4, which combines the unique VerA ITR-adjacent sequence with optimized backbone configuration, was designated the ssAAV MuteFree-VerA vector (**Figure 2B**).

To comply with regulatory standards for GMP manufacturing, the ampicillin resistance gene (Amp) in ssAAV2.4 was subsequently replaced with a kanamycin resistance gene (Kana), generating the ssAAV MuteFree-VerA-Kana variant. Consistently, no ITR mutations were detected in this construct (**Table 1**). We also tested ssAAV2.5, constructed to be identical to ssAAV2.4 except replacing the VerA ITR-adjacent sequence with VerB, which showed equally good ITR stability while preserving structural integrity of the plasmid (**Table 1**). Together, these results showed that ssAAV MuteFree vectors solved ITR instability problems without adding any additional sequences, altering ITR sequences, or requiring special incubation conditions.

### MuteFree system eliminates ITR mutations in scAAV

The baseline scAAV plasmid, scAAV1.0 (**Figure 2A**), contains a 143-nt 5’ ITR and a 113-nt 3’ ITR-Δtrs, each flanked by an 11-nt GC-rich sequence (73% GC) of distinct sequence composition as compared to ssAAV1.0. It carries the same expression cassette between the ITRs as ssAAV1.0. Following the same stability assessment, 31.8% (7/22) of the 5′ ITRs exhibited mutations characterized by the loss of either the B–B′ or C–C′ arm, and one mutation of losing C-C’ arm was detected in the 3′ ITR-Δtrs (**Table 2**). The screening for the ITR-adjacent sequences found two versions of the 11-nt sequence that increased stability of the ITR. One of them, referred to as version 1 (Ver1), has 36% GC. The other one, referred to as version 2 (Ver2), has 27% GC. We next generated the scAAV2.0 backbone by changing the 143-nt 5′ ITR to the 130-nt version used in ssAAV vectors and incorporating the Ver1 ITR-adjacent sequence. The 5’ ITR mutation rate was then lowered to 19.2% (5/26) (**Table 2**). Based on scAAV2.0, a series of vector backbones was constructed and subjected to the ITR stability test. Like the observations for ssAAV, in scAAV2.1 and 2.2 which contained an inserted insulators of 0.5kb and 1kb, respectively, their 5’ ITR mutation rate was reduced to around 8% (**Table 2**). Due to the same considerations about transfection efficiency and safety, we further expanded our screening and identified scAAV2.3, a backbone without any insulator sequence and the same 5’ ITR-adjacent sequence as in scAAV2.0, 2.1 and 2.2, while also replacing the 3’ ITR with the same 11-nt ITR-adjacent sequence as in 5’ ITR. Stability test of scAAV2.3 showed zero mutation in both ITRs (**Table 2**). Thus scAAV2.3 was designated as the scAAV MuteFree-Ver1 vector (**Figure 2D**), and its kanamycin version, scAAV MuteFree-Ver1-Kana, was tested to be equally stable in both ITRs (**Table 2**). Similar results were obtained when replacing the Ver1 11-nt ITR-adjacent sequence in scAAV2.3 with the Ver2 11-nt ITR-adjacent sequence to build scAAV2.4, designated as the scAAV MuteFree-Ver2 vector (**Table 2**).

**Table 2.**
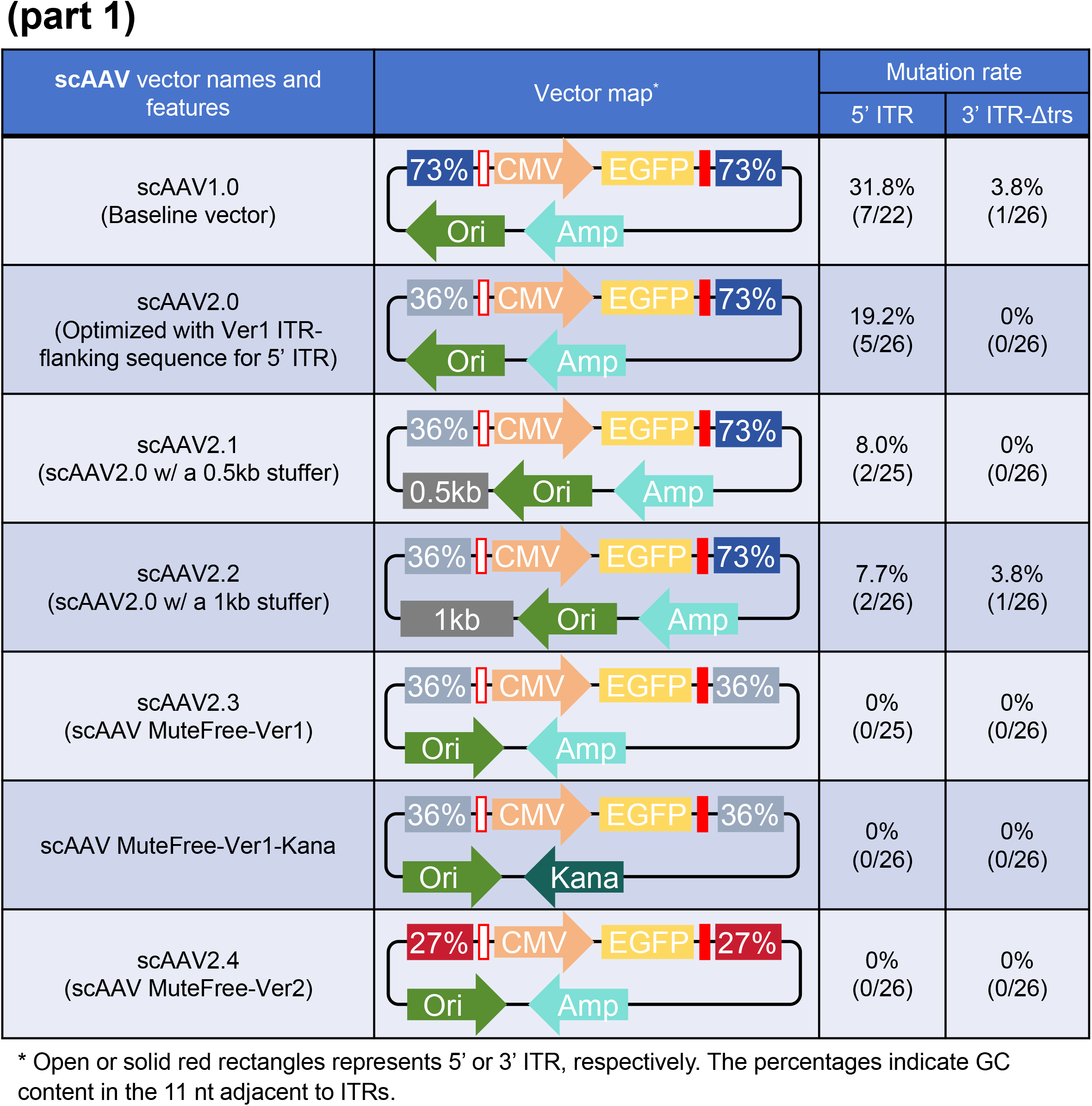
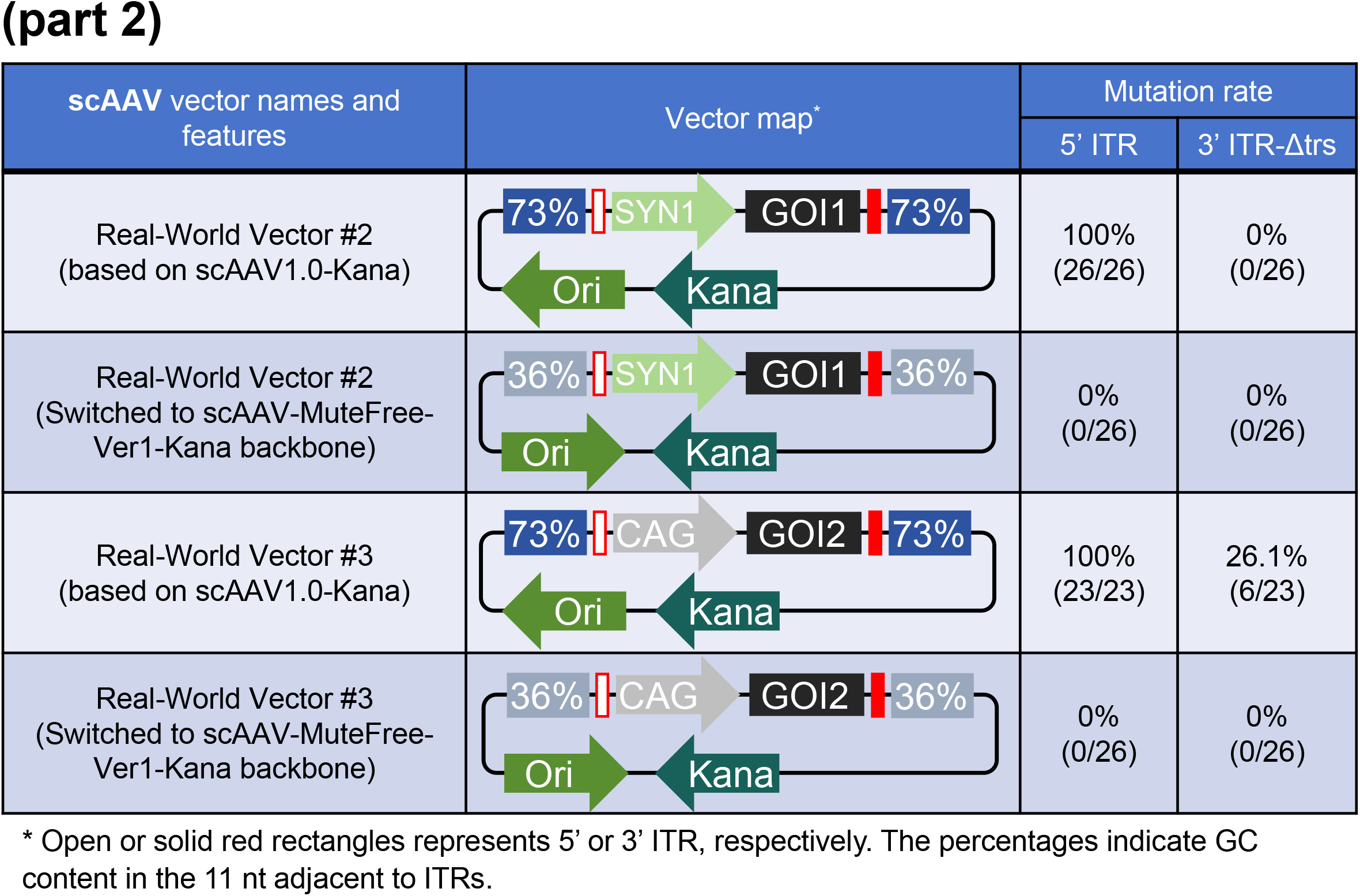
Features of scAAV transfer vectors and their ITR mutation rate.

| scAAV vector names and features | Vector map* | Mutation rate |  |
| --- | --- | --- | --- |
|  |  | 5' ITR | 3' ITR-Δtrs |
| scAAV1.0<br>(Baseline vector) |  | 31.8%<br>(7/22) | 3.8%<br>(1/26) |
| scAAV2.0<br>(Optimized with Ver1 ITR-flanking sequence for 5' ITR) |  | 19.2%<br>(5/26) | 0%<br>(0/26) |
| scAAV2.1<br>(scAAV2.0 w/ a 0.5kb stuffer) |  | 8.0%<br>(2/25) | 0%<br>(0/26) |
| scAAV2.2<br>(scAAV2.0 w/ a 1kb stuffer) |  | 7.7%<br>(2/26) | 3.8%<br>(1/26) |
| scAAV2.3<br>(scAAV MuteFree-Ver1) |  | 0%<br>(0/25) | 0%<br>(0/26) |
| scAAV MuteFree-Ver1-Kana |  | 0%<br>(0/26) | 0%<br>(0/26) |
| scAAV2.4<br>(scAAV MuteFree-Ver2) |  | 0%<br>(0/26) | 0%<br>(0/26) |

\* Open or solid red rectangles represents 5' or 3' ITR, respectively. The percentages indicate GC content in the 11 nt adjacent to ITRs.

Table 2 (part 2)
| scAAV vector names and features | Vector map* | Mutation rate |  |
| --- | --- | --- | --- |
|  |  | 5' ITR | 3' ITR-Δtrs |
| Real-World Vector #2<br>(based on scAAV1.0-Kana) |  | 100%<br>(26/26) | 0%<br>(0/26) |
| Real-World Vector #2<br>(Switched to scAAV-MuteFree-Ver1-Kana backbone) |  | 0%<br>(0/26) | 0%<br>(0/26) |
| Real-World Vector #3<br>(based on scAAV1.0-Kana) |  | 100%<br>(23/23) | 26.1%<br>(6/23) |
| Real-World Vector #3<br>(Switched to scAAV-MuteFree-Ver1-Kana backbone) |  | 0%<br>(0/26) | 0%<br>(0/26) |

\* Open or solid red rectangles represents 5' or 3' ITR, respectively. The percentages indicate GC content in the 11 nt adjacent to ITRs.

ITR stability of the MuteFree AAV vectors was further validated using the conventional restriction enzyme digestion assay. The baseline and MuteFree ssAAV or scAAV vectors were extracted from both the original clones and from cultures after ten serial passages, each performed in triplicate. All plasmid preparations were analyzed by diagnostic restriction digestion using XmaI or AhdI, which cleave intact ITRs within the C–C′ or B–B′ arms, respectively, and are widely used as a rapid assay of ITR integrity. For the baseline ssAAV or scAAV vectors, the intensity of the partially digested bands increased markedly after serial passage (**Figure 3**), suggesting an increasing amount of mutated ITR, which is consistent with single-clone sequencing showing progressive accumulation of ITR mutations (**Table 1** and **2**). In contrast, MuteFree vectors displayed no noticeable digestion products following serial passaging. Together, these results further confirm the markedly improved ITR stability conferred by the MuteFree backbone. Faint bands of partially digested vector were observable with the initial clones of the baseline vectors. As the ITR in those clones have been sequenced validated, we hypothesized that the partial digestion was because high-GC flanking sequence enhanced the ITR secondary structure, making the enzyme recognition sites more inaccessible, instead of actual mutations. Such phenomenon was not observed with the MuteFree vectors, possibly because the modification to the ITR flanking sequence has loosen up the secondary structure, making the restriction enzyme digestion more complete.

**Figure 3.**
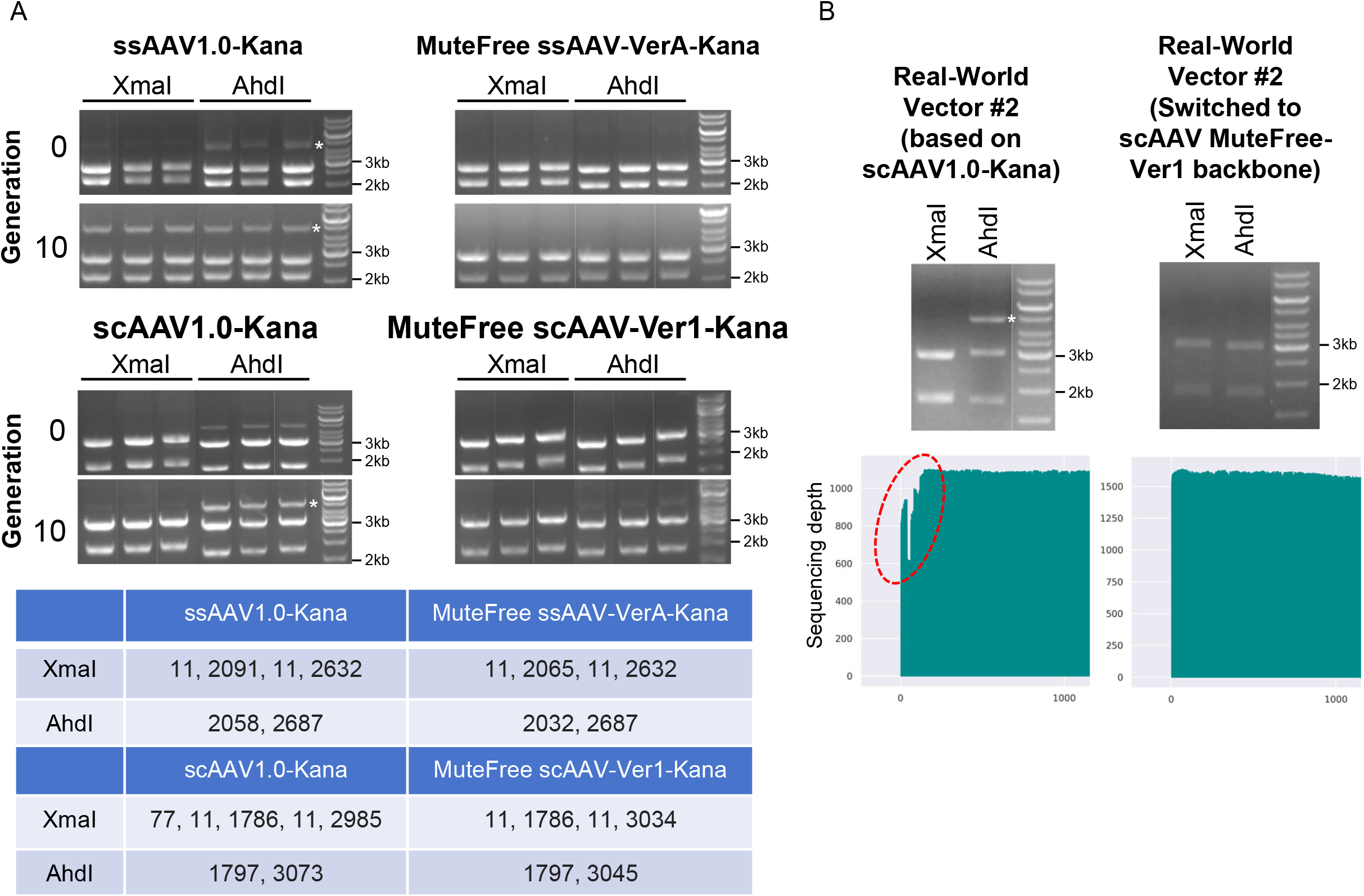
Further stability assessment for ITR on MuteFree vectors. (**A**) The baseline and MuteFree ssAAV or scAAV vectors extracted before and after ten serial passages (generation 0 and 10, respectively) were subjected to diagnostic XmaI or AhdI digestion to determine the integrity of ITR C–C′ or B–B′ arms, respectively. The asterisk indicates the extra band from partial digestion by AhdI due to ITR mutations. The lower table shows the size of the theoretical bands after digestion. (**B**) Real-World Vector #2, which is based on scAAV1.0-Kana backbone (**Table 2**) and its MuteFree counterpart vector were produced using 4L bioreactors and subjected to diagnostic restriction enzyme digestion and Nanopore whole-plasmid sequencing. The red circle in the Nanopore sequencing result highlights significant 5’ ITR mutations detected in Real-World Vector #2 on scAAV1.0-Kana backbone, which is consistent with the presence of the extra band from partial digestion by AhdI due to ITR mutations. The mutations disappeared completely after switching to the MuteFree backbone.

It is important to note that Sanger sequencing of the ITRs can only validate the sequence integrity of the ITRs themselves, but cannot detect structural rearrangements of the plasmids such as large-scale deletions, duplications and inversions, or concatemerization of the plasmids. In contrast, these rearrangements can be detected by restriction digestion. The fact that the aforementioned restriction digestion tests did not uncover any signs of plasmid rearrangements further confirmed the structural stability of the MuteFree system.

Next, we compared the MuteFree vectors and its predecessors in terms of AAV packaging efficiency and the transduction efficiency of the resulting virus. Equal amount of ssAAV or scAAV vectors of the baseline or MuteFree versions were packaged into AAV2 capsid under identical conditions, yielding AAV titers ranging from 2.97-4.24E+13 GC/mL or 4.35-5.46E+13 GC/mL for ssAAV or scAAV vectors, respectively (**Figure 4A**). The full/empty capsid ratios determined using Mass Photometry were within normal range (**Figure 4A**). SDS-PAGE analysis confirmed a consistent ratio of roughly 1:1:10 among the three structural proteins VP1–VP3 across all vectors (**Figure 4B**). Transduction of HEK293 cells at MOI=100 for 48 h resulted in comparable levels of EGFP expression (**Figure 4C**). Collectively, these results demonstrate that modifications to the backbone in ssAAV and scAAV MuteFree vector do not compromise AAV packaging efficiency, capsid composition, and transduction efficiency.

**Figure 4.**
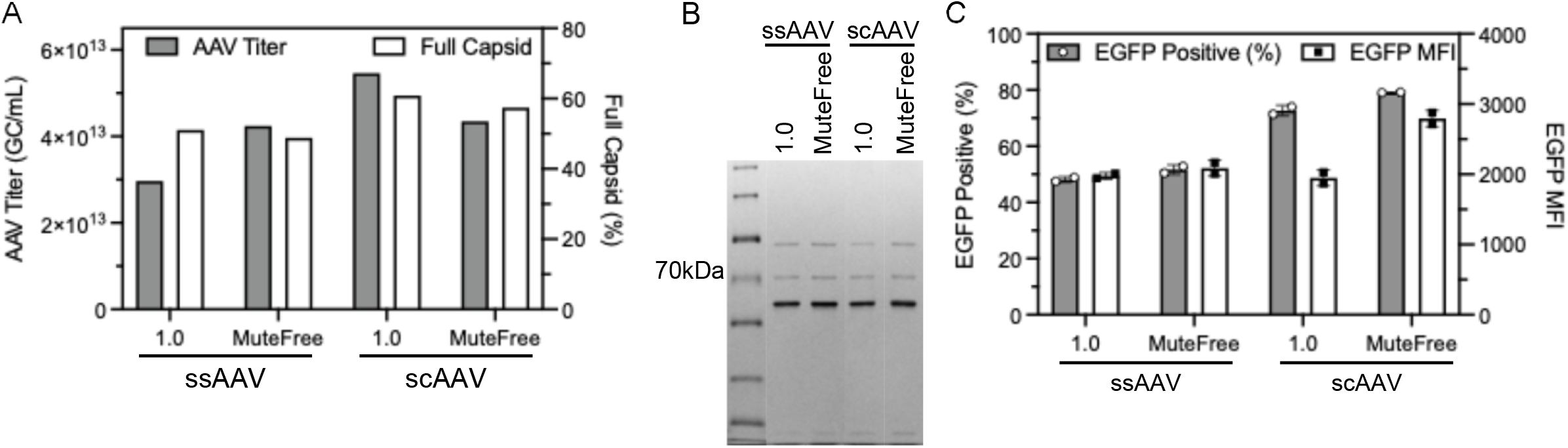
Validation for AAV packaged with MuteFree vectors. Single-stranded (ss) and self-complementary (sc) AAV2 packaged using the 1.0 or MuteFree vector were tested for their (**A**) titer and full capsid ratio, (**B**) purity using silver-stained SDS-PAGE, and (**C**) transduction on HEK293 cells at MOI=100. The expression of AAV-delivered EGFP was assess using flow cytometry at 48 h post transduction. For ssAAV MuteFree vector, the MuteFree-VerA backbone was used. For scAAV MuteFree vector, the MuteFree-Ver1 backbone was used.

### MuteFree system solved instability problems in real-world GMP projects

In our recent experience, ITR instability substantially impeded AAV transfer plasmid production in three GMP manufacturing campaigns, including one ssAAV and two scAAV projects intended for clinical use. These vectors were originally constructed using the ssAAV1.0 or scAAV1.0 backbone. Although clones with intact ITRs were selected at the onset of these projects, 5′ ITR mutations repeatedly emerged during scale-up production. Consistent with these observations, ITR stability testing showed that 50% of the ssAAV and 100% of the scAAV plasmids carried 5′ ITR mutations, predominantly involving loss of the B–B′ or C–C′ arms (**Table 1** and **Table 2**). These projects were originally scheduled for cancellation due to severe ITR instability. But after re-cloning the transfer vectors using the ssAAV or scAAV MuteFree backbone, stability tests demonstrated complete preservation of both 5′ and 3′ ITRs, with no mutations detected in any clone (0/26) (**Table 1** and **2**). The scAAV1.0 and its corresponding MuteFree scAAV vectors of one campaign were produced in 4L bioreactors. The latter showed excellent ITR integrity revealed by diagnostic restriction enzyme digestion and Nanopore sequencing, whereas the former had significant 5’ ITR mutation (**Figure** 3B).

## DISCUSSION

The instability of AAV ITR was first reported in 1982, when the AAV genome was cloned into pBR322 to rescue recombinant AAV (8). Over the following decade, additional studies confirmed that ITR mutations frequently occur when AAV transfer plasmids are propagated in *E. coli*, largely due to the palindromic, GC-rich sequences and the stable secondary structures of the ITR. Meanwhile, several critical ITR elements essential for the AAV life cycle were also elucidated. In particular, the Rep binding element (RBE) within the A–A′ region, the terminal resolution site (trs), and the B–B′/C–C′ arm structures were identified as key features required for viral genome replication. Instability of the ITR can result in the loss of these essential components, reducing AAV yield by more than 10-fold and increasing the non-empty capsid ratio 5–6 fold (13).

Several conventional strategies have been applied to mitigate ITR instability. The most straightforward one is to propagate AAV vectors in *E. coli* strains that are defective in recombinase activity, such as VB UltraStable (*rec*A1), NEB Stable (*rec*A1), and Stbl3 (*recA*13), which effectively reduces recombination-driven mutations, but their effect on the overall ITR stability is limited. Using low copy number plasmid backbone for the AAV transfer vector has also been proven effective (14). Incubating the *E. coli* at a lower temperature (30°C instead of 37°C) for a short time limits the replication of the transfer plasmid, thus reducing the chance of mutation. Both solutions lower ITR mutations at the cost of lowering the yield of AAV transfer plasmid, which is not a concern when producing research-grade AAV as the yield can be offset by larger fermentation volumes. However, when scaling up the transfer plasmid production for a GMP-grade AAV project, larger fermentation volume dramatically increases production cost and process complexity, which adds to the cost, timeline and uncertainty of a GMP project.

Recently, alternative strategies have been proposed. A rationally engineered ITR with reduced GC content (60%) maintained functional encapsulation and gene delivery (15). Although originally designed to reduce CpG-mediated immunogenicity, this modified ITR remained stable after six passages, but yielded only one-third as much virus as the wild-type ITR. Another study combined high-temperature incubation (42°C), which potentially destabilizes ITR hairpins, with an *E. coli* strain deficient in the enzyme responsible for cleaving DNA hairpins (16). This approach enabled production of transfer vectors containing full-length (145 bp) ITRs at a lower mutation rate. But accessibility to the special *E. coli* strain severely limits the application of the method, and the plasmid produced under 42°C was just one fourth that under 37°C.

In this study, we addressed the long-standing issue of ITR instability through systematic optimization to the ITR-adjacent sequences and the plasmid backbone configuration. The synergistic effect of the two approaches made the resulting vectors, designated ssAAV or scAAV MuteFree, exhibit complete ITR stability, with no detectable mutations observed after ten serial passages representing about 166 population doublings, which is a dramatic improvement compared to the 31.8–48.1% ITR mutation rates in the original ssAAV or scAAV vectors (**Tables 1** and **2**). These optimizations did not introduce additional sequences or alter any viral or transgene elements. Furthermore, AAV2 particles produced from the MuteFree vectors displayed comparable yield, purity, and transduction efficiency to those generated using the unmodified vectors (**Figure 4**). Moreover, the application of the ssAAV/scAAV MuteFree vectors rescued three real-world GMP manufacturing projects from cancellation and reduced the 5′ ITR mutation rate from a range of 50-100% to 0% (**Table 1 and 2, Figure 3B**).

Our results demonstrated that ITR stability is largely affected by the flanking sequence. Our data also indicated that the asymmetric instability between the 5′ and 3′ ITRs is possibly due to their proximity to plasmid replication origin in the direction of plasmid replication. This observation is consistent with a recent report showing that shortening the Ori-to-ITR distance from 481 bp to 241 bp increased the proportion of mutant ITRs by approximately 60% (16).

In sum, the MuteFree system presented here effectively addressed ITR instability without altering ITR sequences, *E. coli* fermentation conditions, and the choice of the host strain. This improvement enhanced the robustness of therapeutic AAV vector production while reducing cost and uncertainty in GMP-grade manufacturing for clinical applications.

## METHODS

### Plasmid cloning and validation

The baseline ssAAV (VB231204-1056gvv) and scAAV (VB241109-1146jkd) plasmids were modified using a combination of Gibson assembly and restriction/ligation cloning to generate all the tested variants in the study. Plasmids were chemically transformed to *E. coli* stbl3, and transformants were cultured on LB agar plate supplemented with 100 µg/mL ampicillin or 50 µg/mL kanamycin based on the marker gene on the plasmid at 37°C overnight.

### Bacterial culture

*E. coli* single clones were kept in LB with 25% glycerol at −80°C for long-term preservation. *E. coli* cultures were incubated in LB with corresponding antibiotics based on the plasmid at 37°C and 250 rpm for overnight before plasmid preparation.

### Plasmid validation by sequencing

Plasmids were extracted from overnight culture and validated using a combination of Sanger sequencing for ITR and third-generation sequencing for the whole plasmid.

### Plasmid validation by restriction enzyme digestion

Purified plasmids were digested with restriction enzymes, XmaI or AhdI, from NEB following the vendor’s protocol.

### AAV packaging, titration, and transduction

AAV serotype 2 (AAV2) was packaged in 293T cells using the triple transfection method following optimized VectorBuilder protocols. The titer of AAV was measured by qPCR, and the purity was assessed by subjecting denatured AAV capsids to SDS-PAGE followed by silver staining. To test the transduction potency of AAV2 packaged with different transfer plasmids, viral particles were added to fresh HEK293 cells plated 1 day earlier and reaching 30∼50% confluency. Expression of EGFP was assess by flowcytometry 48 h post transduction.

## ACKNOWLEDGMENTS

This study was supported by VectorBuilder.

